# Next-Generation Sequencing Could be a Promising Diagnostic Approach for Pathogen Detection: Pathogenic Analysis of Pediatric Bacterial Meningitis by Next-Generation Sequencing Technology Directly from Cerebrospinal Fluid Specimens

**DOI:** 10.1101/340273

**Authors:** Ling-yun Guo, Yong-jun Li, Lin-lin Liu, Hong-long Wu, Jia-li Zhou, Ye Zhang, Wen-ya Feng, Liang Zhu, Bing Hu, Hui-li Hu, Tian-ming Chen, Xin Guo, He-ying Chen, Yong-hong Yang, Gang Liu

## Abstract

**Background:** Bacterial meningitis remains one of the major challenges in infectious diseases, leading to sequel in many cases. A prompt diagnosis of the causative microorganism is critical to significantly improve outcome of bacterial meningitis. Although various targeted tests for cerebrospinal fluid (CSF) samples are available, it is a big problem for the identification of etiology of bacterial meningitis.

**Methods:** Here we describe the use of unbiased sequence analyses by next-generation sequencing (NGS) technology for the identification of infectious microorganisms from CSF samples of pediatric bacterial meningitis patients in the Department of Infectious Diseases from Beijing Children’s Hospital.

**Results:** In total, we had 99 bacterial meningitis patients in our study, 55 (55.6%) of these were etiologically confirmed by clinical microbiology methods. Combined with NGS, 68 cases (68.7%) were etiologically confirmed. The main pathogens identified in this study were Streptococcus pneumoniae (n=29), group B streptococcus (n=15), Staphylococcus aureus (n=7), Escherichia coli (n=7). In addition, two cases with cytomegalovirus infection and one with Taenia saginata asiatica were confirmed by NGS.

**Conclusions:** NGS could be a promising alternative diagnostic approach for critically ill patients suffering from bacterial meningitis in pediatric population.

**Summary:** We conducted the study for the identification of microorganisms by next-generation sequencing directly from CSF samples of pediatric bacterial meningitis patients. And the study showed that NGS could be a promising alternative diagnostic approach for bacterial meningitis in pediatric population.

## BACKGROUND

Bacterial meningitis, also known as purulent meningitis, is caused by a variety of gyogenic infections. Although the incidence in infants and children has decreased since the use of conjugated vaccines targeting *Haemophilus influenzae type b* (Hib), *Streptococcus pneumoniae* (*S. pneumoniae*) and *Neisseria meningitides* (*N. meningitides*), bacterial meningitis continues to be an important cause of mortality and morbidity in neonates and children throughout the world^[1, 2]^. The causative pathogens of bacterial meningitis depend on different age of the patient and predisposing factors^[1–3]^.

Pathogen identification is of paramount importance for bacterial meningitis. At present, the pathogen of bacterial meningitis is still mainly based on Gram stain and bacterial culture. However, CSF culture can be negative in children who receive antibiotic treatment prior to CSF examination^[1]^. Because of the limitations of clinical laboratory testing, more than half of the central nervous system infection cases cannot be clearly diagnosed^[4]^. Although non-culture methods including multiplex PCR and latex agglutination, etc. have been used in clinical microbiology^[1]^, only one or several specific pathogens could be targeted by these kinds of technology, let alone rare pathogens.

In recent years, the emergence of powerful NGS technology have enabled unbiased sequencing of biological samples due to its rapid turnaround time^[5]^. Wilson et al^[6]^ presented a case of neuroleptospirosis, resulting in a dramatic clinical improvement with intravenous penicillin after identifying leptospira infection in the CSF by unbiased NGS technology. Unbiased NGS could facilitate identification of all the potential pathogens in a single assay theoretically^[7]^. Herpes simplex virus1, herpes simplex virus 2 and human herpes virus type 3 were detected using NGS technology from four cases with clinically suspected viral meningoencephalitis respectively. And the results were further validated using PCR^[8]^. Further, Yao et al^[9]^ detected *Listeria monocytogenes* in CSF from three patients with meningoencephalitis by NGS. These reports highlight the feasibility of applying NGS of CSF as a diagnostic method for central nervous system (CNS) infection. However, the majority of reports are comprised of single case reports and few studies have been reported in the application of NGS for pathogen detection from CSF samples of bacterial meningitis patients, especially in pediatric populations. In this study, we would like to use the NGS technology to detect directly from the CSF samples of children with bacterial meningitis and evaluate the feasibility and significance of the NGS technique on the pathogenic identification of bacterial meningitis.

## METHODS

### Study Population

Cases of bacterial meningitis in patients of age between 29days and 18 years old from Oct.23^th^ 2014 through Dec.31^th^ 2016 in the Department of Infectious Diseases, Beijing Children’s Hospital were included. Patients with disagreement to collecting CSF, CSF volume < 1ml and bloody CSF were excluded. This tertiary health care hospital is a National Children’s Medical Center with 970beds that treats more than 3 million outpatients and 70,000 hospitalized patients every year. Records for all patients including demographic data, clinical features and laboratory findings were obtained.

### Diagnosis of bacterial meningitis

Any child with sudden onset of fever (> 38.5°C rectal or 38.0°C axillary) and neck stiffness, altered consciousness or other meningeal symptoms were considered to be suspected patients. A case that is laboratory-confirmed [culture method and/ or antigen detection methods (Alere BinaxNow^®^*Streptococcus pneumoniae* Antigen Card)] by identifying a bacterial pathogen from CSF or blood in a child with clinical symptoms consistent with bacterial meningitis is a proven case. Otherwise, a suspected case with CSF examination showing at least one of the following is a probable case: a turbid CSF appearance, leukocytosis (> 100 cells/mm^3^), leukocytosis (10-100 cells/ mm^3^) and either elevated protein (> 100 mg/dl) or decreased glucose level (< 40 mg/dl). These criteria are consistent with the World Health Organization (WHO) case definition^[10]^.

### Sample collection

CSF was collected in accordance with standard procedures, snap frozen and stored at −80° C in Bio bank for Diseases in Children Beijing Children’s Hospital, Capital Medical University.

### DNA extraction, library construction, and sequencing

DNA was extracted directly from the clinical samples using the TIANamp Micro DNA Kit (DP316, Tiangen Biotech, Beijing, China). DNA libraries were constructed through end-repaired adapter added overnight, and by applying polymerase chain reaction amplification to the extracted DNA. Quality control was carried out using a bioanalyzer (Agilent 2100, Agilent Technologies, Santa Clara, CA, USA) combined with PCR to measure the adapters before sequencing. DNA sequencing was then performed using the BGISEQ-100 platform (BGI-Tianjin, Tianjin, China)^[11]^.

### Data treatment and analysis

High-quality sequencing data were generated after filtering out low-quality, low-complexity, and shorter reads. To eliminate the effect of the human sequences, the data mapped to the human reference genome (hg19) were excluded using a powerful alignment tool called Burrows-Wheeler Alignment^[12]^. The remaining data were then aligned to the Microbial Genome Database, which includes bacteria, viruses, fungi, and protozoa. Finally, the mapped data were processed by removing duplicate reads for advanced data analysis. A no redundant database that included all the published genomes of microorganisms was downloaded from the National Center for Biotechnology Information (ftp://ftp.ncbi.nlm.nih.gov/genomes/). The depth and coverage of each species were calculated using Soap Coverage from the SOAP website (http://soap.genomics.org.cn/).

### Ethics Statement

This study was reviewed and approved by the Ethics Committee of Beijing Children’s Hospital Affiliated to Capital Medical University (No.2017-74). Written informed consents were obtained from all patients or their legal surrogates.

### Statistical Analysis

Categorical variables were compared using the chi-squared test or Fischer’s exact test, as appropriate. Continuous variables within two groups were compared using the independent t-test for parametric data and the Mann-Whitney U test for non-parametric data. *P* values <0.05 were considered statistically significant. All statistical analyses were conducted using SPSS 17.0(SPSS Inc. USA).

## RESULTS

### Demographics

A total of 99 cases were finally included in our study. Among them, 68 (68.7%) cases were male, 31 (31.3%) females, and a male to female ratio of 2.2: 1. The median age of onset was 5.7 months. These cases were classified into four age groups: 73 (73.8%) patients aged 29 days to 1 year, 1-year-old to 2 years old group (n = 13, 13.1%); 2 years old to 5 years old group (n = 4, 4.0%); greater than 5 years old group (n = 9, 10.0%). 57.6% (57 cases) of children were from rural areas. The median time to diagnose bacterial meningitis was 3 (2-7) days.

### Bacterial pathogen detection by clinical microbiology methods

55 cases (55.6 %) had positive clinical pathogen detections. Among them, 33 (33.3%) cases were positive from the CSF culture, 32 (32.3%) cases were positive from the blood culture, and 13 (13.1%) cases of them were positive both from CSF and blood culture. 21 (21.2 %) cases had positive results of Alere BinaxNow^®^ *Streptococcus pneumoniae* Antigen test. In terms of pathogen distribution, *S. pneumoniae* infection was the primary pathogen, 23 (41.8 %) cases of children with the presence of this bacterial infection. Followed by *Streptococcus agalactia* in 11 cases (20.0 %), *Escherichia coli* in six cases (10.9 %), *Staphylococcus aureus* in five cases (9.09 %), *Listeria monocytogenes* in four cases (7.3 %). There was also one case of *Enterococcus faecalis, Streptococcus bacteria, Streptococcus mitis, Haemophilus influenzae, Streptococcus bovis*, and *Eickhelium echinobacterium*, respectively.

### NGS results

#### NGS overall data

99 samples of cerebrospinal fluid were detected by NGS, and generated 2, 852,780,707 raw reads. After the removal the short read length (length ≤ 35bp), the low complexity and low-quality reads, 2,846,199,682 were obtained. The average number of raw reads and clean reads for each sample were 28,815,967 and 28,749,492, respectively. The mean rate of 95.35% mapped to the human genome. And finally, the average ratios of the reads mapped to bacterial, viral and fungal genome were 0.0216%, 0.000015% and 0.000405%, respectively.

### Determination of pathogens by NGS technology

A total of 10 kinds of pathogenic microorganisms were determined by NGS. There were six kinds of Gram-positive bacteria, two kinds of Gram-negative bacteria, one kind of virus (two cases of cytomegaloviruses) and one kind of parasite (*Taenia saginata*) among them. Among the pathogenic bacteria, *S. pneumoniaewas* still the predominant pathogen accounting for 51.2% of bacteria, followed by *Streptococcus agalactiae in 19.5% of bacteria, Staphylococcus aureus* in 7.3%, *Enterococcusfaecali in 7.3%, Escherichia coli* in 4.9%, *Listeria monocytogenes* in 4.9%. The coverage, depth and unique reads of different spices were between 0.0046% −68%, 1-2, 2-67550, respectively. (Table 1 and Figure 1).

**Figure 1.**
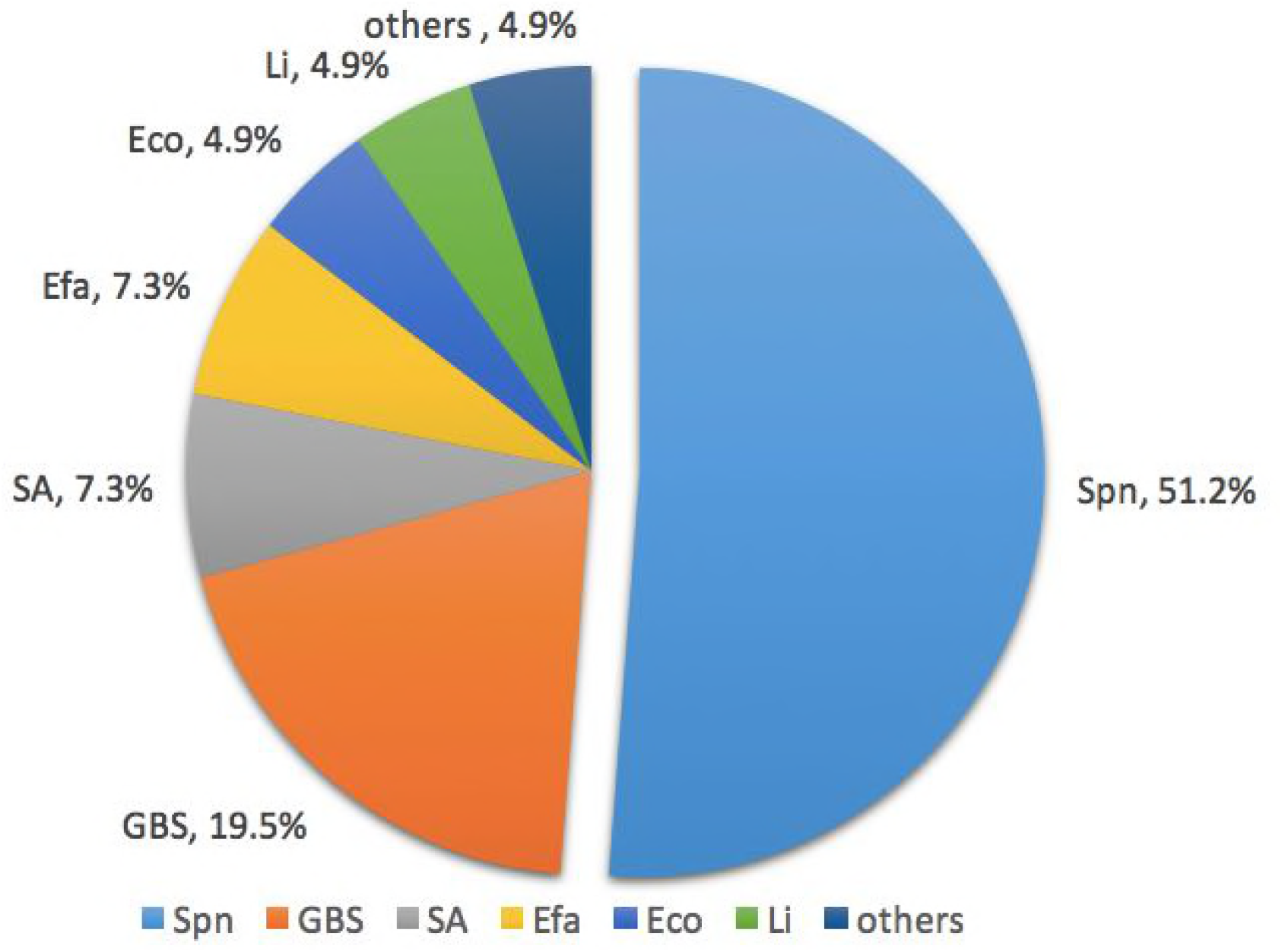
The distribution of different bacterial pathogens by NGS. Note: Spn, *Streptococcus pneumoniae*; GBS, *group B Streptococcus*; SA, *Staphylococcus aureus*; Eco, *Escherichia coli*; Li, *Listeria monocytogenes*; Efa, *Enterococcus faecium*; others includes *Pseudomonas aeruginosa* and *Staphylococcus pasteuri*.

**Table 1.** Pathogenic microorganisms determined by NGS

| Type | Pathogens | No. | Coverage | Depth | Unique Reads |
| --- | --- | --- | --- | --- | --- |
| Gram-positive<br>bacteria | *Streptococcus pneumoniae* | 21 | 0.011%-11% | 1-1.1 | 4-1105 |
|  | *Streptococcus agalactiae* | 8 | 0.0046%-25% | 1-1.2 | 2-12004 |
|  | *Staphylococcus aureus* | 3 | 0.26%-2.4% | 1 | 56-1380 |
|  | *Enterococcus faecium* | 3 | 0.03%-0.74% | 1 | 17-394 |
|  | *Listeria monocytogenes* | 2 | 0.04%-5% | 1 | 8-2561 |
|  | *Staphylococcus pasteurii* | 1 | 20% | 1.1 | 5716 |
| Gram-negative<br>bacteria | *Escherichia coli* | 2 | 0.34%-0.46% | 1 | 32-65 |
|  | *Pseudomonas aeruginosa* | 1 | 68% | 2 | 67550 |
| Virus | *Cytomegalovirus* | 2 | 0.45%-1.1% | 1 | 20-49 |
| Parasite | *Taenia saginata asiatica* | 1 | 0.58% | 1 | 9610 |
| Total |  | 44 | 0.0046%-68% | 1-2 | 2-67550 |

### Combination of NGS and clinical microbiology methods

Combined with clinical pathogen detection methods and NGS technology, 76 kinds of microorganisms were determined while 44 of them were detected by NGS and 55of them were detected by clinical microbiology methods. A total of 68 cases (68.7%) had positive pathogens, of which 67 cases (67.7%) had positive bacterial pathogen results. NGS increased the positive rate of pathogen detection by 13.1% (the positive rate of pathogen detection increased from 55.6% to 68.7%) and 23.6% of the pathogens were independently detected by the NGS method.

In terms of analyzing specific pathogen, a total of 29 *S. pneumoniae* were detected by combination methods, while six of them were independently detected by the NGS method; a total of 15 *S. agalactiae* were detected by combination methods, while four of them were independently detected by the NGS method; a total of seven *Staphylococcus aureus* were detected by combination methods, and two of them were independently detected by the NGS method; a total of seven Escherichia coli were detected by combination methods, and one of them were independently detected by the NGS method(Figure 2).Overall, *S. pneumoniae* (n=29, 39.7%), *S.agalactiae* (n=15, 20.5%), *Staphylococcus aureus* (n=7, 9.6%), and *Escherichia coli* (n=7, 9.6%) were the most frequently detected pathogens in this study. In addition, there were four cases of *Listeria monocytogenes* (5.5%), three of *Enterococcus faecium* (4.1 %) (Figure 3).

**Figure 2.**
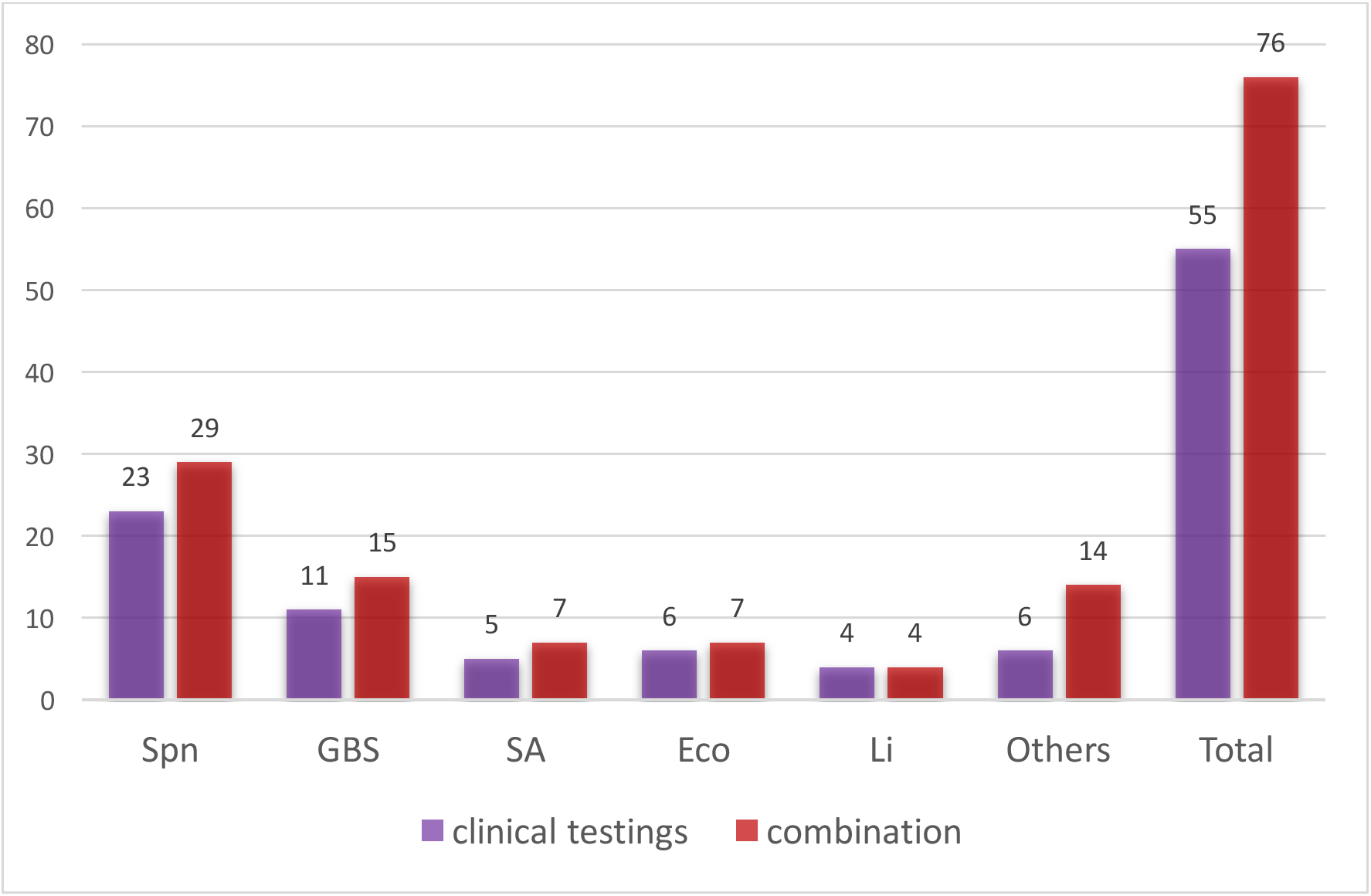
Number of different pathogens detected by clinical testings and combination with NGS. Note: Spn, *Streptococcus pneumoniae*; GBS, *group B Streptococcus*; SA, *Staphylococcus aureus*; Eco, *Escherichia coli*; Li, *Listeria monocytogenes*; others includes *Enterococcus faecalis, Enterococcus faecalis, Streptococcus pyogenes, Streptococcus mitis, Streptococcus bovis, Haemophilus influenza, Streptococcus bovis, Eikenella corrodens, Pseudomonas aeruginosa* and *Staphylococcus pasteuri, Cytomegalovirus and Taenia saginata asiatica*

**Figure 3.**
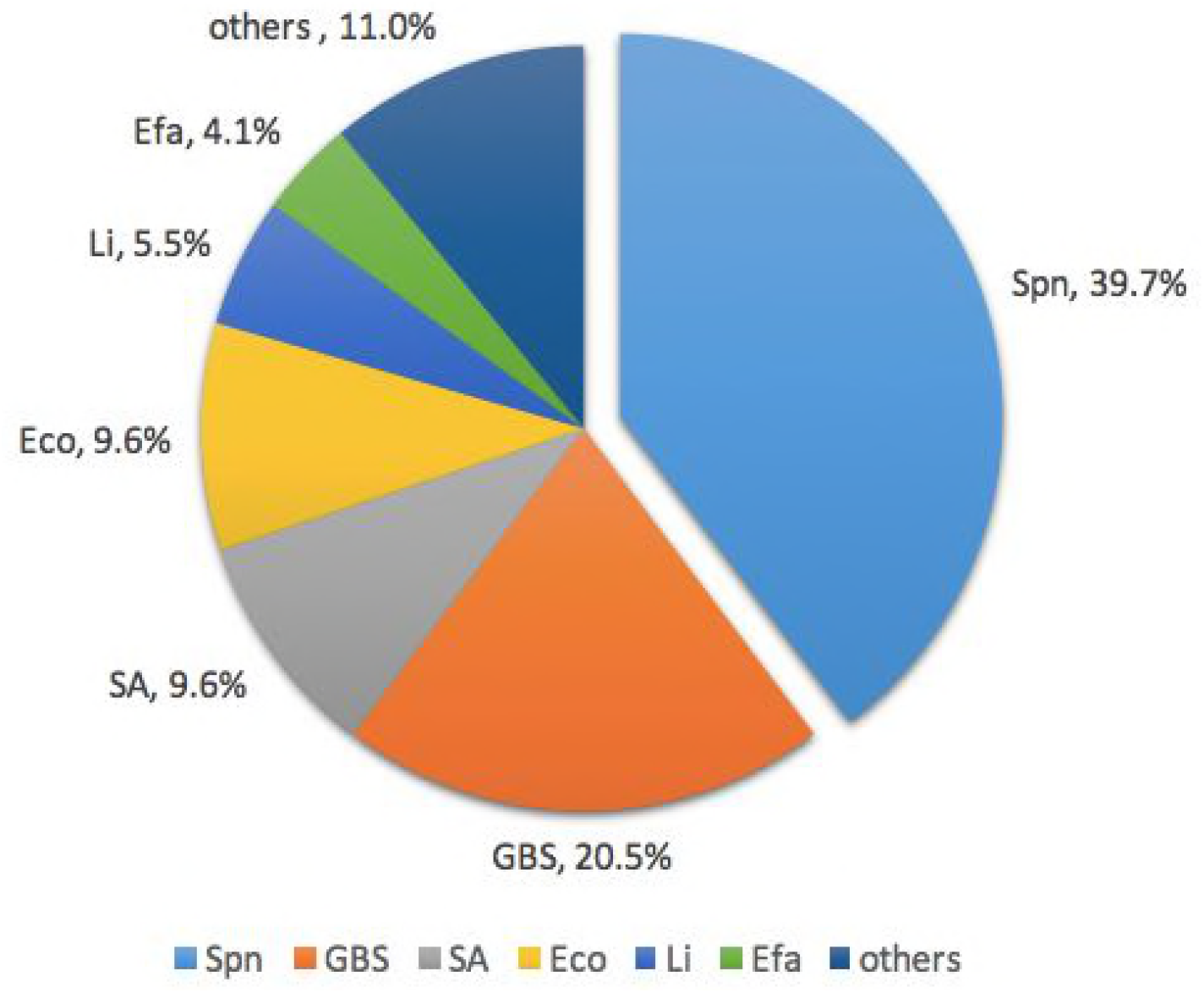
The distribution of bacterial pathogens by combination of clinical methods and NGS. Note: Spn, *Streptococcus pneumoniae*; GBS, *group B Streptococcus*; SA, *Staphylococcus aureus*; Eco, *Escherichia coli*; Li, *Listeria monocytogenes*; *Efa, Enterococcus faecium*; others including *Enterococcus faecalis, Streptococcus pyogenes,Streptococcus mitis, Haemophilus influenza,Streptococcus bovis, Eikenella corrodens, Pseudomonas aeruginosaand Staphylococcus pasteuri;others includesEnterococcus faecalis, Streptococcus pyogenes,Streptococcus mitis, Haemophilus influenza,Streptococcus bovis, Eikenella corrodens, Pseudomonas aeruginosa*and *Staphylococcus pasteuri*

### Comparison of the NGS-positive and NGS-negative groups

The NGS-positive group comprised 41 (41.4%) patients who had positive bacterial results by NGS. The NGS-negative group comprised 58 cases. Patients in the NGS-positive group had generally shorter days in terms of duration from onset to the sample collection (median days 15 vs. 31 days, *P* = 0) and shorter onset days than those in the pathogen-negative group (median days 6 vs. 25 days, *P* = 0.005). With regard to blood inflammatory indices, patients in the NGS-positive group were frequently found to have higher neutrophils (66.7% vs. 54.7%, *P*= 0.002) than those in the NGS-negative group. The two groups of patients did not differ significantly in age or diagnosis duration. In addition, there was no significant difference in the simultaneously CSF WBC count between members of these two groups, and they had CSF similar protein and glucose levels (Table 2).

**Table 2.** Comparison of NGS-positive and NGS-negative groups^a^

| Items |  | NGS-positive <sup>b</sup><br>(n=41) | NGS-negative<br>(n=58) | *P* |
| --- | --- | --- | --- | --- |
| Age (days) |  | 230(69.5-425) | 116.5(65.5-392.5) | 0.379 |
| Duration from onset to the sample collection<br>(days) |  | 15(8.5-28) | 31(17-45) | 0 |
| Onset (days) |  | 6(4-21) | 25(4-41) | 0.005 |
| Diagnosis Duration (days) |  | 4(2-8) | 3(1-6.75) | 0.66 |
| Peripheral | WBC($\times 10^9/L$) | 17.3±11.9 | 13.5±7.3 | 0.074 |
| Blood | neutrophils (%) | 66.7±15.7 | 54.7±20.2 | 0.002 |
| | hemoglobin (g/L) | 104.0±19.9 | 103.8±17.2 | 0.943 |
|  | C-reactive protein (mg/L) | 29 (11.0-101.1) | 27.7(8-108.5) | 0.393 |
| 1<sup>st</sup> CSF | WBC ($\times 10^6/L$) | 910(336-2825) | 403(98-2040) | 0.125 |
|  | multinuclear cells (%) | 70(38.75-85) | 60(22-75) | 0.165 |
|  | monocyte (%) | 24.3(15-55.5) | 38.5(23.0-75.0) | 0.099 |
|  | protein (mg/L) | 1278(837-2234) | 1340(596-1982) | 0.485 |
|  | glucose (mmol/L) | 1.47(0.87-2.74) | 2.48(1.02-2.48) | 0.12 |
| Simultaneously | WBC ($\times 10^6/L$) | 10(2-17) | 11(2-37) | 0.432 |
| CSF <sup>c</sup> | multinuclear cells (%) | 10(4-16) | 12(5-20) | 0.723 |
|  | monocyte (%) | 20(12-28) | 30(15-54) | 0.051 |
|  | protein (mg/L) | 557(296-1149) | 594(367-1079) | 0.85 |
|  | glucose (mmol/L) | 2.85(2.43-3.46) | 2.90(2.46-3.66) | 0.869 |

NGS, next generation sequencing; WBC, white blood cell; CRP, C-reactive protein; CSF, cerebrospinal fluid.
<sup>a</sup> Results are reported as the median (interquartile range), or as the percentage.
<sup>b</sup> NGS-positive group refers to the cases who had positive bacterial NGS results.
<sup>c</sup> Simultaneously CSF refers to the CSF results by clinical testing at which time CSF is detected by NGS.

## Discussion

It can be seen from our data that NGS can improve the overall positive rate of pathogen detection, especially in clinically negative samples. The results of clinical pathogen detection and NGS pathogen showed that 68 cases (68.7%) were positive in 99 cases of bacterial meningitis, 67 cases (67.7%) were positive for bacterial pathogens. Further analysis of data, we found that 13 cases had positive pathogens in the group of clinical microbiological-negative group (44 cases). NGS increased the positive rate of pathogen detection by 13.1% (the positive rate of pathogen detection increased from 55.6% to 68.7%). Although there are few results in terms of NGS using in the large population of bacterial meningitis, there is increasing evidence of a role for NGS in the work-up of undiagnosed encephalitis and meningitis^[13]^. Brown JR et al recommend NGS should be considered as a front-line diagnostic test in chronic and recurring encephalitis. And the results in our study consistent with a study of NGS which conducted in patients with sepsis, and detectability were significantly increased from 12.82% to 30.77%^[14]^. It indicates that an NGS-based approach has great potential to detect causative pathogens of infectious diseases.

Unbiased NGS technology amplifies all nucleic acids present in a clinical sample, including both host and microbes without requiring primers for targeted amplification, and can potentially generate microbial sequence data offering the capability of identifying a variety of organisms-bacteria, virus, fungi, or parasite^[15]^. As can be seen from our study, 95.35% of the NGS sequencing data were compared to the human genome, whereas less than <1 % to the pathogen genome. Previous studies referred to the application of NGS in clinical pathogen detection showed similar situation in both bacterial and/or virus infections^[14]^. In addition, this method clearly shows the advantages of recognizing multi-pathogens infection. Alexander and other researchers have emphasized that it is important to rich the genome database in order to enhance the completeness analysis of the results. If you only add a part of gene sequence to the database, this may reduce the sensitivity of the assay^[16,17]^. Therefore, in order to give full play to the potential of NGS, it is necessary to set up a well suitable local database and coverage pathogen genome as much as possible. The results in our study showed that the primary pathogen was still *S. pneumoniae*. 29 cases of *S. pneumoniae* were detected, accounting for 39.7% of the total bacterial pathogens. It is worth noting that the second place of the pathogen is *S. agalactiae* in 15 cases, accounting for 20.8% of the total detected bacterial pathogens. This is consistent with the rising trend of *S. agalactiae*, which has been reported in our previous study^[18]^. Furthermore, we found two cases with cytomegaloviruses infection and one patient with *Taenia saginata*. These three patients were diagnosed as bacterial meningitis definitely, thus NGS clearly shows the advantages of recognizing multi-pathogens infection. However, the role of other microorganisms, e.g. cytomegaloviruses and *Taenia saginata*, in the process of bacterial meningitis should be further discussed in the future.

The interpretation of the NGS results is of great important and difficult in some situations. NGS pathogen detection usually obtains a variety of suspected pathogens gene sequences, and how to distinguish between pathogens and contaminating “background bacteria” becomes critical. Procedures including clinical specimen collection, sample preservation, transportation and the experimental procedures, may cause pollution and lead to contaminate the specimen mixed with so called “background” microbial gene sequence, which may affect the NGS data analysis^[15,19,20]^, resulting in false positive results and hinder the doctors. Similar to previous reports, we found that the most common background bacterium was *Propionibacterium acnes* in our study (data not shown here)^[14]^.

We compared NGS-positive and NGS-negative groups, and analyzed factors that could affect NGS results. Patients in the NGS-positive group had generally shorter days in terms of duration from onset to the sample collection (median days 15 vs. 31 days, *P* = 0) and shorter onset days than those in the pathogen-negative group (median days 6 vs. 25 days, *P* = 0.005). In terms of time between two groups, we found that 32 (94.12%) of the samples were less than 42 days (data did not show) in the NGS-positive group. Only 2 patients received cerebrospinal fluid samples > 42 days. It indicated that the earlier detected the more positive possibility. In addition, it is noting that there was no significant difference in the simultaneously CSF WBC count between two groups, and they had CSF similar protein and glucose levels. These altogether suggests that the original severity of inflammation, not the inflammation status while the simultaneously NGS detects might determine the NGS results. The impact of time on the results of NGS may be related to the nucleic acid content in the specimen, but this is still to be further validated by future clinical studies of large samples.

To our knowledge, the present study is one of the largest case series of using NGS in (pediatric) bacterial meningitis patients around world. However, this study has some limitations. It was a single center, hospital-based retrospective design, which introduces the possibility of unrecognized biases, incomplete data collection. Thus, further prospective multi-center studies are necessary to enrich the generalizability of NGS technology using in pediatric bacterial meningitis.

In summary, our study demonstrated the comprehensive diagnostic ability of NGS in identifying etiology of bacterial meningitis and the factors associated with the NGS results. As a new technology of detection, NGS could be a promising alternative diagnostic powerful tool for pathogen detection allowing us to detect organisms in critically ill patients suffering from bacterial meningitis.

## Conclusions

NGS could be a promising alternative diagnostic approach for critically ill patients suffering from bacterial meningitis in pediatric population.

## Acknowledgements

We would like to thank Professor Hongzhi Guan at the Department of Neurology, Peking Union Medical College Hospital for his assistance and valuable suggestions of this study. we would like to thank Bio bank for Diseases in Children Beijing Children’s Hospital, Capital Medical University for their contribution of storing the CSF samples with standard procedures. In addition, we would like to thank Xue Yao at BGI-Shenzhen for her coordination in this study.

## Author contributors

All of the authors had access to the full dataset (including the statistical reports and tables) and take responsibility for the integrity of the data and the accuracy of the data analysis. GLY, LYJ, YYH and LG conceived and designed the study. GLY, LLL, ZY, FWY, ZL, HB, HHL, CHY, CTM, GX were involved in case and sample collection, analysis, and interpretation of the data. GLY, LYJ, WHL, ZJL participated in the laboratory analysis of CSF. GLY wrote the first draft of the paper. GLY, YYH, and LG reviewed and approved the final report.

